# RAPPID: a platform of ratiometric bioluminescent sensors for homogeneous immunoassays

**DOI:** 10.1101/2020.10.31.363044

**Authors:** Yan Ni, Bas J.H.M. Rosier, Eva A. van Aalen, Eva T.L. Hanckmann, Lieuwe Biewenga, Anna-Maria Makri Pistikou, Bart Timmermans, Chris Vu, Sophie Roos, Remco Arts, Wentao Li, Tom F.A. de Greef, Frank J. M. van Kuppeveld, Berend-Jan Bosch, Maarten Merkx

## Abstract

Heterogeneous immunoassays such as ELISA have become indispensable in modern bioanalysis, yet translation into easy-to-use point-of-care assays is hindered by their dependence on external calibration and multiple washing and incubation steps. Here, we introduce RAPPID (<u>Ra</u>tiometric <u>P</u>lug-and-<u>P</u>lay Immuno<u>d</u>iagnostics), a “mix-and-measure” homogeneous immunoassay platform that combines highly specific antibody-based detection with a ratiometric bioluminescent readout that can be detected using a basic digital camera. The concept entails analyte-induced complementation of split NanoLuc luciferase fragments, photoconjugated to an antibody sandwich pair via protein G adapters. We also introduce the use of a calibrator luciferase that provides a robust ratiometric signal, allowing direct in-sample calibration and quantitative measurements in complex media such as blood plasma. We developed RAPPID sensors that allow low-picomolar detection of several protein biomarkers, anti-drug antibodies, therapeutic antibodies, and both SARS-CoV-2 spike protein and anti-SARS-CoV-2 antibodies. RAPPID combines ratiometric bioluminescent detection with antibody-based target recognition into an easy-to-implement standardized workflow, and therefore represents an attractive, fast, and low-cost alternative to traditional immunoassays, both in an academic setting and in clinical laboratories for point-of-care applications.

## Introduction

The widely recognized need for more personalized, patient-centered healthcare has urged the development of cheap, easy-to-use, yet reliable biomolecular diagnostics. Such technology would not only accelerate biomolecular research in academic and clinical environments, but also allow for fast diagnostic decision making when access to clinical laboratories is unavailable or cost-prohibitive. Affinity-based immunoassays, such as ELISA and other heterogeneous immunoassays, form the cornerstone of today’s clinical bioanalytical toolbox and offer both the specificity required to minimize crosstalk and the modularity to allow application to a broad range of clinically relevant analytes. Translation of these immunoassays into point-of-care applications has proven challenging, due to the requirement of multiple washing and incubation steps, external calibration and time-consuming optimization of sensor surface immobilization and assay conditions^1^. Lateral flow immunoassays (LFIAs) have been introduced as cheap and relatively easy-to-use point-of-care tests, but they suffer from limited sensitivity and are primarily used as qualitative tests^2,3^.

Single-step affinity-based detection assays performed directly in solution would not only speed up traditional laboratory immunoassays, but would also be particularly attractive for point-of-care testing outside of the laboratory setting by non-expert users^4,5^. Several promising optical biosensor platforms have been reported that translate analyte binding directly into a conformational change, resulting in a change of signal intensity or color^6,7^. Of these, sensors based on bioluminescence are particularly suited for measurements directly in complex media such as blood with minimal sample handling, as they do not require external excitation and therefore do not suffer from issues related to autofluorescence or light scattering^8,9^. The Johnsson lab pioneered the development of semisynthetic sensor proteins based on bioluminescence resonance energy transfer (BRET) for the detection of small-molecule drugs such as methotrexate and digoxin, and important metabolites such as NAD^+^ and NADPH^10–13^. Alternatively, the LUMABS platform was developed to enable detection of specific antibodies directly in blood plasma with the use of a smartphone camera^14–17^. Both sensor platforms employ ratiometric bioluminescent detection, which allows quantitative measurements and enables integration in paper- and thread-based analytical devices for low-cost point-of-care detection^11,18,19^. Although LUCID and LUMABS have a modular architecture, successful sensor development requires the availability of suitable ligand-binding domains and ligand analogs, or specific epitope peptides or mini-protein domains^20,21^, respectively. Moreover, while extensive protein engineering has yielded sensors with a greater than 10-fold change in emission ratio^13^, the dynamic range of BRET sensors is typically 2-4 fold^14,17^.

Here, we present RAPPID (<u>Ra</u>tiometric <u>P</u>lug-and-<u>P</u>lay Immuno<u>d</u>iagnostics) as a new, comprehensive platform of ratiometric bioluminescent sensors that can be readily adopted to suit a broad range of biomolecular targets. RAPPID is based on the complementation of antibody-conjugated split NanoLuc luciferases^22,23^ to detect the formation of a sandwich immunocomplex in solution with a high intrinsic signal-to-background output (**Fig. 1a**). Although methods for the generation of antibody-split luciferase fusion proteins are available, they are based on non-specific conjugation or cumbersome recombinant expression of antibody fusions^13,24^. RAPPID does not require additional protein engineering and uses protein G-mediated photoconjugation to generate well-defined antibody conjugates directly from commercially available antibodies^25,26^. Additionally, we introduce a green light-emitting calibrator luciferase as a simple method to provide a robust blue-over-green ratiometric readout and direct internal calibration. The ratiometric signal of RAPPID sensors is stable over time and less sensitive to experimental conditions and substrate concentration, enabling us to effectively address this fundamental problem of intensiometric bioluminescent assays.

**Figure 1.**
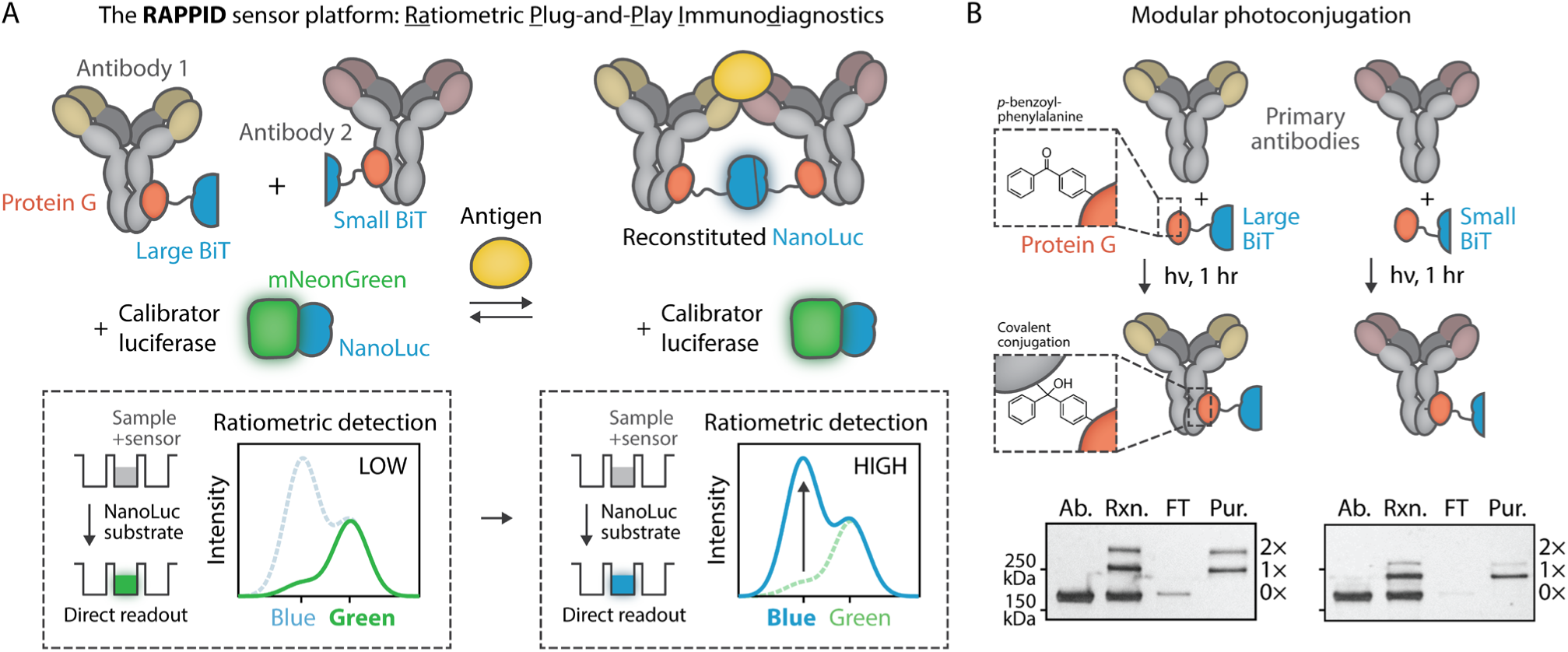
General overview and design elements of the RAPPID sensor platform. **a**, Schematic representation of the concept of antigen detection using a RAPPID sensor. A pair of primary antibodies is functionalized with a split version of the NanoLuc luciferase (Large BiT and Small BiT, respectively) through use of protein G-based photoconjugation. Antigen binding induces co-localization of the antibody pair and subsequent reconstitution of NanoLuc, resulting in the emission of blue light upon conversion of substrate (furimazine; *λ*_*max*_∼460 nm). Ratiometric detection is achieved by addition of a calibrator luciferase, consisting of a tight fusion of NanoLuc to fluorescent acceptor mNeonGreen, which produces a green signal (*λ*_*max*_∼520 nm) upon substrate conversion through bioluminescent resonance energy transfer (BRET). The homogeneous solution-based assay is conducted by mixing the sample and sensor components with the NanoLuc substrate and recording the emission spectrum. In the absence of antigen (left) no split complementation occurs resulting in predominantly green emission from the calibrator luciferase (low blue/green ratio), while the presence of antigen (right) results in NanoLuc reconstitution and an increase in blue-light emission (high blue/green ratio). **b**, Schematic overview of the antibody conjugation method. Fusion between the adapter protein G carrying the photoreactive non-natural amino acid *p*-benzoylphenylalanine and either Large BiT or Small BiT (Gx-LB and Gx-SB, respectively) enables covalent coupling to the constant Fc-binding domain of most IgG-type antibodies, including all human IgGs, by exposure to long-wavelength UV light (*λ*=365 nm). Non-reducing SDS-PAGE analysis of the photoconjugation reaction (Rxn.) illustrates successful coupling of primary antibody (Ab., 10 µM) to either one (indicated with 1×) or two (2×) protein G adapters (10 µM). Reactions were performed in PBS (pH 7.4) for 1 hr on ice. Subsequent small-scale Ni^2+^-affinity chromatography removed unconjugated antibody (FT, flow through) resulting in pure antibody conjugates (Pur.).

Development and implementation of a RAPPID assay for a new biomolecular target is straightforward and consists of three steps: 1) the selection of a pair of (commercial) antibodies that bind the target analyte, 2) crosslinking of the antibodies to the protein G-luciferase adapters using a simple one-hour illumination protocol, and 3) addition of both antibody-luciferase conjugates, the calibrator luciferase and the Nanoluc substrate to the sample, followed by detection of the emission ratio of blue over green light (**Fig. 1a**). The ratiometric luminescent signal can be recorded with a standard digital camera, obviating the need for additional high-tech, expensive equipment. Here, the broad scope and excellent analytical performance of RAPPID is demonstrated by developing assays for a range of clinically relevant biomarkers, including cardiac troponin I, C-reactive protein, three anti-drug-antibodies, and two therapeutic antibodies, displaying robust, up to 36-fold increases in emission ratio. Finally, we demonstrate that the modularity of the RAPPID workflow enables prompt development of novel immunoassays for emerging target analytes such as SARS-CoV-2 spike proteins and anti-SARS-CoV-2 antibodies.

## Results

### Sensor design and characterization

To establish a universally applicable sensing platform, we employed protein G-based photoconjugation for the synthesis of sensor components directly from commercially available primary antibodies. Both the 18-kDa Large BiT (LB) and the 1.3-kDa Small BiT (SB) fragment of split NanoLuc were genetically fused via a semi-rigid peptide linker to a protein G adapter domain (Gx), carrying the photocrosslinkable non-natural amino acid *p*-benzoylphenylalanine (Gx-LB and Gx-SB, respectively, see **Fig. 1b** and **Supplementary Fig. 5**). Expression was performed in *Escherichia coli* with amber codon suppression using an engineered orthogonal aminoacyl tRNA synthetase/tRNA pair^27,28^, which afforded Gx-LB and Gx-SB in high yields after affinity chromatography purification (respectively ∼18 mg/L and ∼5 mg/L, see **Methods** and **Supplementary Fig. 6**). Next, antibody conjugation was performed by illuminating reaction mixtures with long-wavelength UV light (*λ*=365 nm) in phosphate-buffered saline (pH 7.4) for 1 hr on ice, covalently crosslinking Gx-LB and Gx-SB to the Fc-domain of the antibody. SDS-PAGE analysis revealed a mixture of mono-conjugated, bi-conjugated and non-conjugated antibodies (**Fig. 1b**). Although the conjugation efficiency can be further improved by increasing the relative concentration of Gx-LB and Gx-SB, we chose to use an equal molar ratio of protein to antibody to minimize the presence of non-conjugated Gx-LB and Gx-SB, which could contribute to non-specific background complementation. If necessary, non-conjugated antibodies can be removed by small-scale Ni^2+^-affinity chromatography, targeting the N-terminal polyhistidine tag of Gx-LB and Gx-SB (**Fig. 1b**). Since protein G binds to the majority of IgG-type antibodies, including all human and many other mammalian isotypes, this approach offers a general method for the rapid synthesis of antibody-luciferase conjugates from commercially available antibodies.

To establish proof of principle and characterize the performance of the RAPPID sensor platform, we first developed a sensor targeting cardiac troponin I (cTnI), an important protein biomarker for myocardial injury that requires highly sensitive detection at picomolar concentrations (**Fig. 2a**)^29^. To this end, Gx-LB and Gx-SB were photoconjugated to a pair of commercially available anti-cTnI antibodies with distinct binding epitopes (19C7 (mIgG2b) and 4C2 (mIgG2a), respectively, **Fig. 1b**). Subsequently, increasing concentrations of cTnI were added to a reaction mixture consisting of 1 nM of both antibody-luciferase components and incubated for 30 min at room temperature followed by the addition of NanoLuc substrate. A maximal 281-fold increase in luminescence intensity was observed upon addition of cTnI, with a detection regime spanning four orders of magnitude and a limit of detection of 4.2 pM (**Fig. 2b** and **Supplementary Table 1**). The luminescent output follows a bell-shaped dose-response curve, as evidenced by a decrease in signal at cTnI concentrations >10 nM (**Fig. 2b**). This ‘hook’ effect is well-known and occurs because at high target concentrations formation of inactive binary complexes is favored over the formation of the enzymatically active ternary complex^30,31^. These initial results demonstrate that target binding effectively stimulates NanoLuc complementation, resulting in a sensitive bioluminescent signal output with a high signal-to-background ratio.

**Figure 2.**
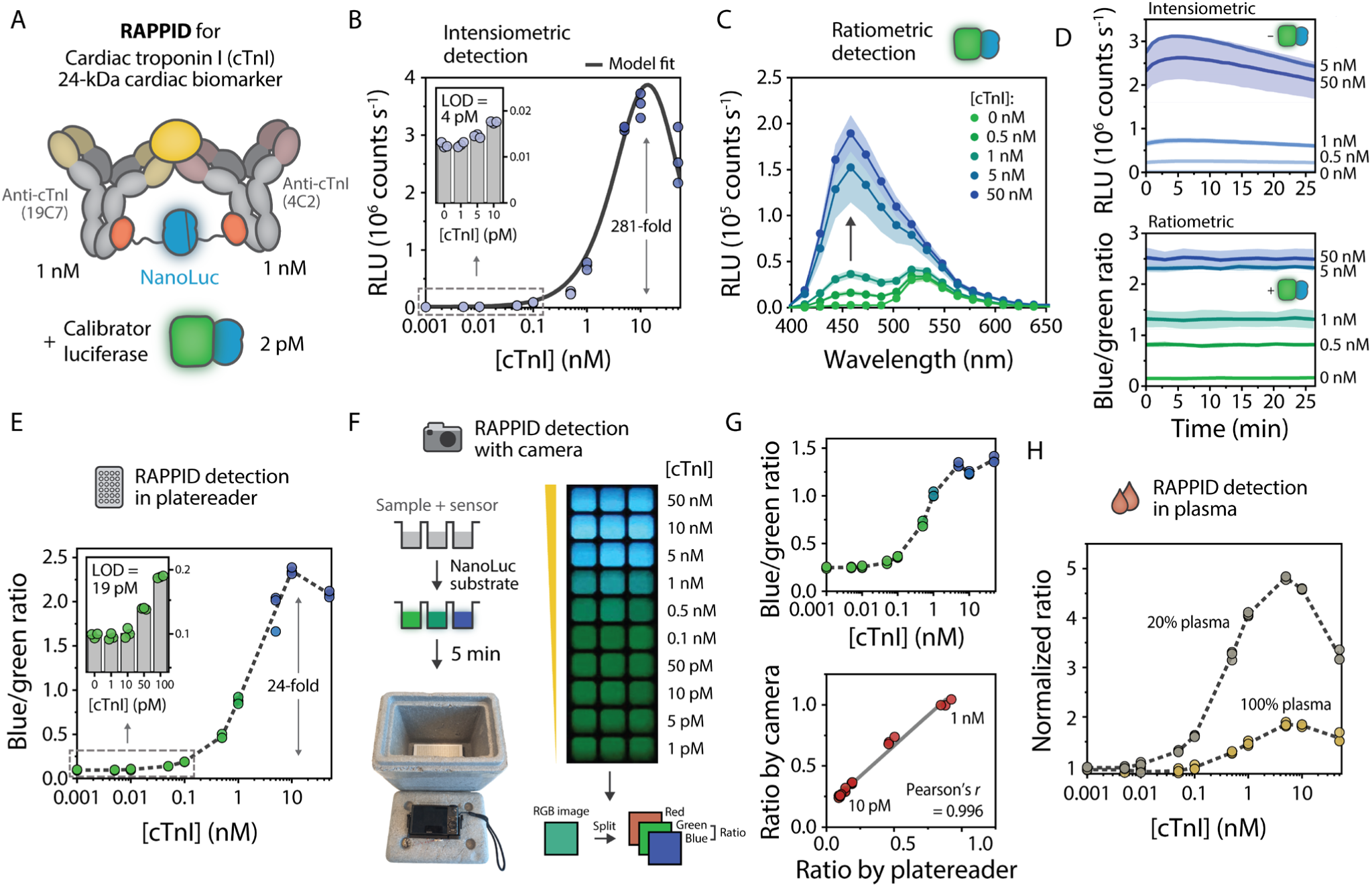
RAPPID enables robust ratiometric detection of cardiac troponin I (cTnI) in buffer and plasma and allows detection with a digital camera. **a**, Schematic illustration of the sensor components used for the detection of cTnI. All experiments were performed in triplicate in buffer (PBS (pH 7.4), 0.1% (w/v) BSA), with 1 nM anti-cTnI (19C7) conjugated to Gx-LB, 1 nM anti-cTnI (4C2) conjugated to Gx-SB, and 2 pM calibrator luciferase, unless indicated otherwise. Reaction mixtures were incubated for 1 hr at room temperature prior to addition of NanoLuc substrate and recording of the emission spectra. The blue/green ratio for ratiometric detection was calculated by dividing bioluminescent emission at 458 nm by emission at 518 nm. **b**, Intensiometric sensor response curve for cTnI. RLU, relative luminescence units. Inset shows sensor response at low target concentrations. Limit of detection (LOD) was determined at 4.2 pM with a maximal fold change of 281. An overview of all sensor characteristics including LOD confidence intervals can be found in **Supplementary Table 1**. Experimental data was fitted to a thermodynamic model (solid line, see **Supporting Figure 4**). **c**, Bioluminescence spectra at various concentrations of cTnI in the presence of 2 pM calibrator luciferase. **d**, Sensor output over time for the experiments described in **b** (top, intensiometric) and **c** (bottom, ratiometric). Data in **c**,**d** are represented as mean ± s.d. **e**, Sensor response curve for increasing concentrations of cTnI. Blue/green ratios were calculated from the spectra in **c**, and show a 24-fold maximal increase in signal. The inset displays sensor behavior in the low-concentration regime as indicated by the dashed rectangle, with a limit of detection (LOD) of 19 pM. **f**, Bioluminescent emission does not require external excitation and therefore enables detection with a conventional digital camera (Sony DSC-RX100) inside a box (left bottom). Photograph of the sensor response for increasing cTnI (right), taken 5 min after addition of the NanoLuc substrate. **g**, Response curve calculated from the photograph in **f** (top) and the correlation between results from the plate reader and digital camera (bottom) for cTnI concentrations below 1 nM. Solid line indicates linear fit. Correlation analysis yielded a Pearson correlation coefficient (*r*) = 0.996. **h**, Sensor response in 100% (yellow) and 20% (gray, diluted in buffer) human blood plasma, recorded in the plate reader. In **e**,**g**,**h** individual data points are represented as circles, bars represent mean values, and dashed lines connect mean values.

To provide a means for understanding and tuning the response characteristics of the RAPPID sensor, we developed a thermodynamic model that analyzes the equilibria involved in target-induced formation of the luminescent ternary complex (**Supplementary Fig. 1** and **Supplementary Fig. 2**). We investigated critical parameters that can be used to modulate the sensor response and performed nonlinear fitting of the experimental data to the model (**Fig. 2b**).. The model predicts that sensor sensitivity and response are strongly dependent on the antibody affinities and that the detection regime can be tuned by varying the concentrations of antibody-luciferase conjugates (**Supplementary Fig. 3a, 3c**). We confirmed the latter predictions by performing RAPPID assays at sensor component concentrations of 10, 1, and 0.1 nM respectively (**Supplementary Fig. 8**). An additional parameter that affects sensor performance is the intrinsic affinity between split NanoLuc components LB and SB. As predicted by the model, attenuating the interaction between LB and SB from *K*_*d*_ = 2.5 µM to *K*_*d*_ = 190 µM does not allow effective luciferase complementation in the sandwich complex. Experimentally, increasing the strength of the interaction (*K*_*d*_ = 0.28 µM) does not enhance luciferase complementation much further in the presence of the analyte, but does results in a higher background signal in the absence of analyte and therefore a smaller dynamic range (**Supplementary Fig. 3b**). These observations indicate that the thermodynamic model can inform the behavior of RAPPID sensors and facilitate rational design of their molecular characteristics.

**Figure 3.**
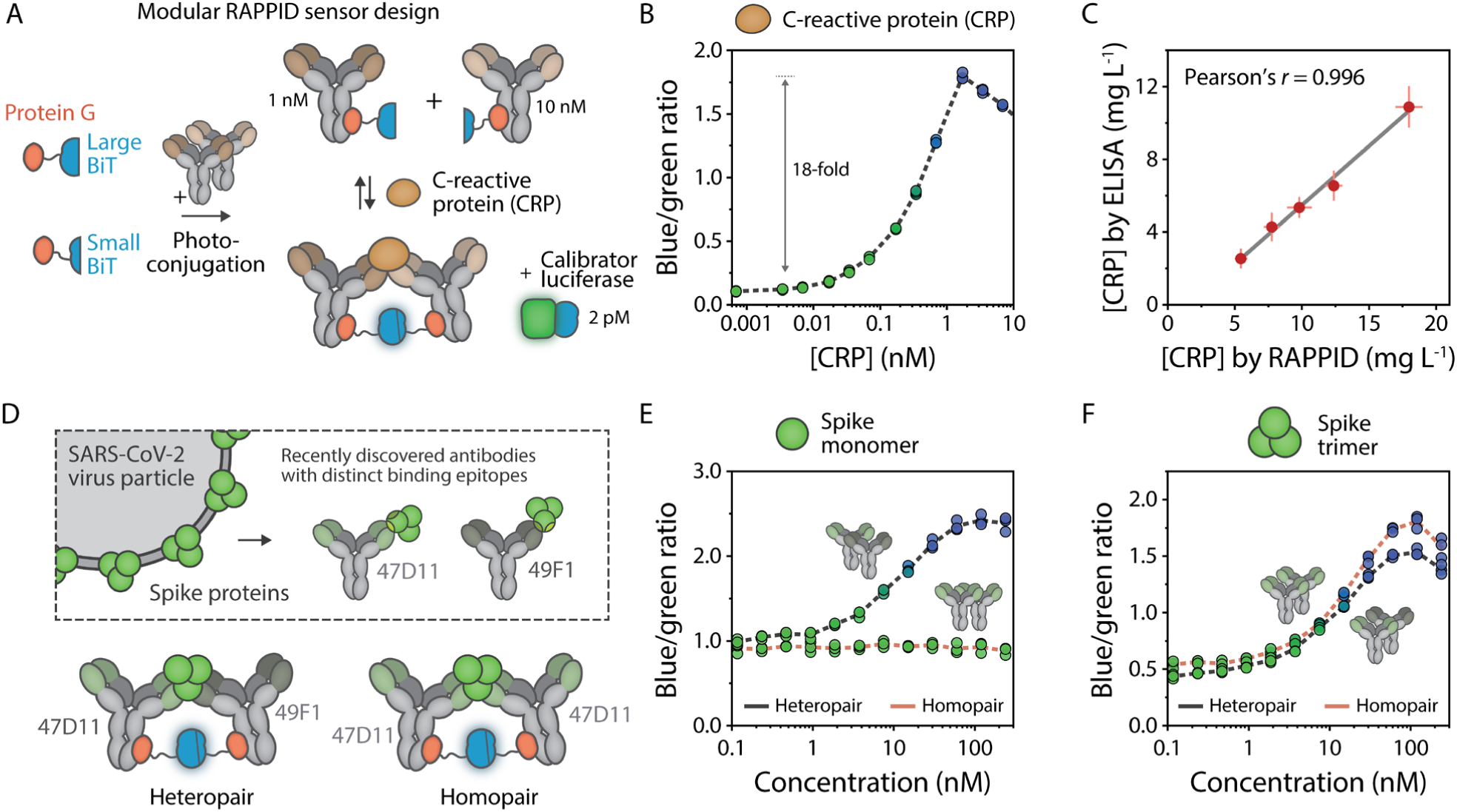
RAPPID sensors for detection of C-reactive protein (CRP) and SARS-CoV-2 spike proteins. **a**, Schematic illustration of the sensor components used for the detection of CRP. All experiments were performed in triplicate, with 1 nM anti-CRP (C6) conjugated to Gx-LB, 10 nM anti-CRP (C135) conjugated to Gx-SB, and 2 pM calibrator luciferase. Reaction mixtures were prepared in buffer (PBS (pH 7.4), 0.1% (w/v) BSA) and incubated for 30 min at room temperature prior to addition of NanoLuc substrate and recording of the emission spectra. The blue/green ratio was calculated by dividing emission at 458 nm by emission at 518 nm. **b**, Sensor response curve for increasing concentrations of CRP, displaying an 18-fold maximal increase in signal. **c**, Correlation of CRP concentrations measured by ELISA and RAPPID. Samples were prepared in diluted blood plasma and quantified using calibration curves obtained in buffer (see **Supplementary Fig. 13**). Error bars represent mean ± s.d. (*n* = 4). Solid line indicates linear fit. Regression analysis yielded a Pearson’s correlation coefficient (*r*) = 0.996. **d**, Schematic overview of RAPPID sensors for the detection of trimeric SARS-CoV-2 spike proteins. Recently discovered antibodies^43,44^ targeting distinct binding epitopes of the immunogenic spike protein were used to generate sensors in either a heteropair or homopair setup. **e**,**f**, RAPPID response curves in response to increasing concentrations of monomeric spike protein (**e**) and native trimeric spike protein (**f**) using both the heteropair (gray) and homopair (red) configuration. Photoconjugations were performed as described in **Supplementary Fig. 14**, and conjugation mixtures were used without further purification. A 200-fold excess of cetuximab was added to suppress background luminescence. Experiments were prepared and performed as described in **a**, but with 1 nM anti-spike (47D11) conjugated to Gx-LB and 10 nM anti-spike (49F1 for heteropair, 47D11 for homopair) conjugated to Gx-SB. In **b**,**e**,**f**, individual data points are represented as circles and dashed lines connect mean values.

An inherent property of intensiometric luciferase assays is depletion of the substrate over time and a concurrent decrease in absolute signal intensity, which hampers reproducibility and prohibits quantitative measurements. This is especially problematic for point-of-care applications, where it is often difficult to calibrate the assay and compensate for variations in substrate concentration, as well as environmental conditions such as pH, temperature and ionic strength. We hypothesized that a calibrator luciferase based on NanoLuc, but with a shifted emission maximum, could be added to the sample together with the sensor components to provide a direct internal calibration by ratiometrically comparing two different emission wavelengths. To construct the calibrator luciferase, we fused NanoLuc to the green fluorescent protein mNeonGreen (mNG-NL), resulting in green luminescent emission (*λ*_*max*_∼520 nm) due to efficient BRET from NanoLuc to the fluorescent acceptor mNeonGreen^32^. The calibrator was added to the cTnI sensor mixture at a final concentration of 2 pM, after which luminescent spectra were recorded at various cTnI concentrations (**Fig. 2c**). In contrast to intensiometric detection, where the luminescent signal quickly reached a maximum and then decayed, the ratiometric signal was stable over a prolonged period of time (**Fig. 2d**). A similar dose response was observed, with a maximal signal at 10 nM cTnI and a limit of detection of approximately 19 pM (compare **Fig. 2b** and **2e**). Although the dynamic range is lower than in the intensiometric assay due to spectral overlap between blue and green emission, the >25-fold change in emission ratio is greater than those typically seen in BRET-based sensor platforms^10,15^. In addition, the calibrator concentration can be adjusted in order to obtain an optimal response at the desired detection regime of the analyte (**Supplementary Fig. 10**).

We also applied the ratiometric nature of the RAPPID sensor to facilitate straightforward signal recording with a digital camera and used the high intrinsic dynamic range to perform measurements directly in blood plasma. Using identical reaction conditions as described for **Fig. 2c**, the cTnI concentration-dependent color change from blue to green was clearly observed in images recorded with a standard digital camera (**Fig. 2f**). The emission ratios were calculated from the blue- and green-color channels of the RGB image and showed a similar dose response as in the plate reader assay, with a linear correlation for cTnI concentrations ≤1 nM (**Fig. 2e** and **2g**). Additionally, we assessed sensor performance in human blood plasma. The luminescent output signal was substantially suppressed in 100% plasma, which can be partially attributed to absorption of blue light by hemoglobin and biliverdin (**Fig. 2h** and **Supplementary Fig. 11**)^33,34^. Nevertheless, the high intrinsic sensitivity of the RAPPID sensor still enabled ratiometric detection, with a reliable 5-fold change in emission ratio in 20% blood plasma (**Fig. 2h**). These results establish the RAPPID sensor platform as a robust ratiometric detection method with a large intrinsic dynamic range. As a result, one-step in-solution measurements can be performed in clinically relevant media such as blood plasma, while facile signal recording with a digital camera should enable rapid diagnostic testing at the point of care.

### Modularity and versatility of RAPPID

To demonstrate that the RAPPID workflow can be implemented for alternative targets, we next developed a sensor for C-reactive protein (CRP), a pentameric protein with clinical relevance as an inflammatory marker in cardiovascular disease^35^, COPD^36^ and other diseases (**Fig. 3a**). Two anti-CRP antibodies (C135 and C6) were photoconjugated to Gx-LB or Gx-SB, respectively, using similar conditions as described for cTnI, and SDS-PAGE was used to confirm efficient conjugation and purification (**Supplementary Fig. 12**). Using 1 nM of C6-LB, 10 nM C135-SB and 2 pM calibrator luciferase, an 18-fold increase in emission ratio was observed when varying the CRP concentration in the pM–nM regime, with an LOD of 2.9 pM (**Fig. 3b**). We then benchmarked the analytical performance of RAPPID to ELISA, which is the current golden standard in hospitals and diagnostic centers for detecting CRP in cardiovascular risk assessment at low mg L^-1^ levels in human blood plasma (approximately 8– 80 nM)^37,38^. A linear correlation (Pearson’s *r* = 0.9906) was observed between CRP concentrations as determined by ELISA and RAPPID (**Fig. 3c** and **Supplementary Fig. 13**), confirming the accuracy of our CRP sensor. The high sensitivity of RAPPID allowed a 50-fold dilution of plasma samples, thereby mitigating possible interference from matrix effects. In addition, the “mix-and-measure” workflow eliminates the need for multiple wash steps and thus substantially reduces total assay time.

After validating the modularity of RAPPID, we sought to demonstrate that our platform is suited for the rapid development of sensors for newly emerging targets. Containment of the current COVID-19 pandemic, caused by the SARS-CoV-2 virus, urgently requires strategies for fast, low-cost, point-of-care viral detection, as no effective drug or vaccine is yet available^39–42^. Two recently reported monoclonal antibodies that target the immunogenic spike glycoprotein of the SARS-CoV-2 virus allowed us to develop RAPPID sensors for viral antigen detection. The antibodies (47D11 and 49F1) bind distinct epitopes on the spike protein which natively exists as a trimer. We constructed two sensor variants, using either both antibodies (heteropair, 47D11-LB with 49F1-SB) or only a single antibody (homopair, 47D11-LB with 47D11-SB) and directly utilized them for sensing the spike protein without further purification (**Fig. 3d** and **Supplementary Fig. 14**). To scavenge non-conjugated Gx-LB and Gx-SB and consequently reduce the background signal in absence of the analyte, we added an excess of non-binding IgG to the assay solution. We hypothesized that we could differentiate between the monomeric and trimeric state of the spike protein, the latter being more likely to represent the active virus, by comparing the heteropair and homopair sensor response. Indeed, only the heteropair sensor variant, in which the antibody-luciferase conjugates are able to bind to distinct epitopes, exhibited a dose response to monomeric spike protein, while no response was observed when using the homopair (**Fig. 3e**). Both sensor variants showed a concentration-dependent increase in emission ratio in response to the spike protein trimer, with a detection regime ranging between 1 nM and 100 nM (**Fig. 3f**). Model simulations suggest that the sensitivity could be further improved by using antibodies with higher affinities (**Supplementary Fig. 3c**). Indeed, when performing similar tests using commercially available anti-spike antibodies with higher affinities, the LOD could be decreased by almost an order of magnitude, between 0.2 and 0.3 nM (**Supplementary Fig. 15** and **Supplementary Fig. 16, Supplementary Table 1**). Taken together, these results illustrate the potential of RAPPID for on-demand development of new biosensors as soon as antibodies become available, and the ability of the platform to distinguish between oligomeric states of proteins.

### Antibody detection using RAPPID

The RAPPID platform also provides new assay formats for the detection of clinically relevant antibodies. Our previously developed LUMABS BRET sensor platform allows sensitive and quantitative detection of specific antibodies, but requires the genetic incorporation of well-defined, high affinity epitopes or mimitopes, which are not always available^14^. Here, we used the RAPPID platform to detect clinically relevant antibodies with complex, discontinuous epitopes and for the detection of heterogeneous antibody responses. We first developed sensors for anti-drug-antibodies (ADAs), which are raised by the immune system against therapeutic antibodies upon repeated administration and are a main reason for rapid clearance^45^. We utilized our photoconjugation strategy to directly couple both Gx-LB and Gx-SB to three important clinical therapeutic antibodies: cetuximab which targets EGFR and is used for colorectal cancer treatment^46^, and adalimumab and infliximab which target TNFα and are prescribed for a variety of autoimmune diseases^47^ (**Fig. 4a** and **Supplementary Fig. 17**). Similar dose-response curves were obtained for all three ADAs (**Fig. 4b**). Because both antibody-luciferase conjugates contain the same antibody that binds to the bivalent ADA target, a statistical mixture containing a fraction of non-luminescent ternary complexes is expected to form. Despite this, the RAPPID sensors exhibited a robust change in emission of 5–6-fold, and LODs between 59–81 pM (**Supplementary Table 1**).

**Figure 4.**
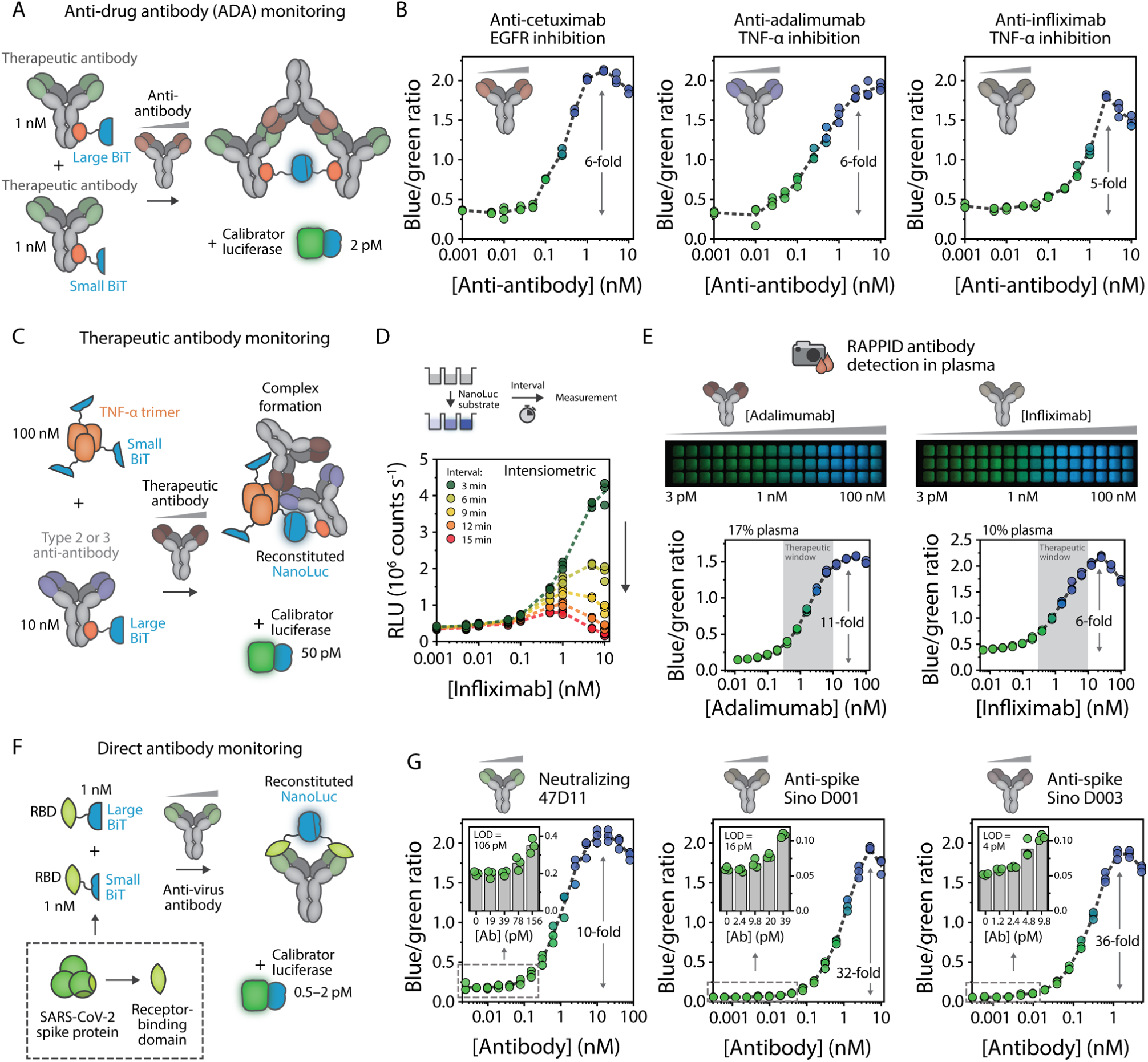
RAPPID enables versatile monitoring of clinically relevant antibodies. **a**, Schematic overview of RAPPID sensors for the detection of anti-drug antibodies (ADAs). Use of a commercially available therapeutic antibody conjugated to both Gx-LB and Gx-SB enables binding of the ADA and formation of a luminescent ternary complex. **b**, Ratiometric sensor response curves of RAPPID sensors for anti-cetuximab, anti-adalimumab, and anti-infliximab, respectively. **c**, Schematic overview of RAPPID sensors for detection of therapeutic antibodies adalimumab and infliximab using target antigen TNFα fused to SB, and the anti-antibody conjugated to Gx-LB. **d**, Intensiometric detection of infliximab without calibrator luciferase. Luminescent measurements were performed at various intervals following addition of the NanoLuc substrate. **e**, Sensor response for adalimumab (left) and infliximab (right) detection in diluted plasma. The same sample was used for detection with a digital camera (top) and the plate reader (bottom). **f**, Schematic overview of RAPPID sensors for direct monitoring of anti-virus antibodies against the receptor binding domain (RBD) of SARS-CoV-2 spike protein. LB and SB were expressed as fusion proteins with the RBD, enabling detection of all RBD-targeting antibodies. **g**, Ratiometric sensor response curves for neutralizing antibody 47D11 (left)^44^, and commercially available anti-spike D001 (center) and D003 (right)^43^. Insets show sensor response at low target concentrations, which allowed calculation of the limit of detection, as indicated in the graphs. Experiments in **b**,**d**,**e**,**g** were performed in triplicate at sensor and calibrator concentrations as indicated in **a**,**c**,**f**, respectively. Reaction mixtures were prepared in buffer (PBS (pH 7.4), 0.1% (w/v) BSA) and incubated for 1 hr at room temperature before addition of NanoLuc substrate and recording of the emission spectra. The blue/green ratio was calculated by dividing bioluminescent emission at 458 nm by emission at 518 nm. The fold increase is indicated in each graph. Individual data points are represented as circles, while dashed lines connect mean values. Bars in histograms represent mean values.

Next, we applied RAPPID to detect antibodies that bind structurally complex, discontinuous epitopes, employing it to monitor therapeutic antibodies that target the TNFα homotrimer as a proof of concept^47^. TNFα-binding therapeutic antibodies, such as adalimumab and infliximab, are widely administered anti-inflammatory drugs for the treatment of diseases like rheumatoid arthritis^48^ and inflammatory bowel disease^49^. Therapeutic drug monitoring (TDM) of these antibodies is important in the clinic to achieve optimal treatment efficiencies and to reduce possible side effects. To overcome the issue of creating non-productive complexes as described above for the ADA sensors, we adapted the RAPPID format by introducing a TNFα-SB fusion protein and a Gx-LB-photoconjugated type-2 anti-antibody, binding to the therapeutic antibody, or a type-3 anti-antibody, binding to the complex of the target therapeutic antibody and TNFα (**Fig. 4c** and **Supplementary Fig. 18**). Because of the relatively high therapeutic window of adalimumab and infliximab^50,51^, assays were performed using higher sensor concentrations (10 nM anti-antibody-LB and 100 nM TNFα-SB). As a result, we observed high substate turnover and a quick decrease in luminescence intensity within a few minutes after adding the NanoLuc substrate (**Fig. 4d**). Addition of the calibrator luciferase was therefore critical to allow accurate, time-independent antibody quantification. Robust changes in emission ratio in the therapeutically relevant concentration range for both adalimumab and infliximab were detected directly in 10–17% blood plasma using either a plate reader or a digital camera (**Fig. 4e**), demonstrating the ability of RAPPID sensors to accurately measure concentrations of antibodies in complex media.

The ongoing pandemic caused by SARS-CoV-2 warrants for the development of serological assays that can be used as a tool to evaluate the extent of viral infection in the population and assess the efficacy of potential vaccine candidates. As such, we next aimed to develop a RAPPID variant for the detection of antibodies against the SARS-CoV-2 virus^52,53^. To enable the detection of a wide panel of anti-SARS-CoV-2 antibodies, we adapted the RAPPID sensor by introducing the immunogenic receptor binding domain (RBD) of the SARS-CoV-2 spike protein^54^, fused to either SB or LB (**Fig. 4f**). The bivalent nature of antibodies promotes the complementation of LB with SB, hence producing a bioluminescent signal upon RBD-LB and RBD-SB binding. After successful expression of RBD-LB and RBD-SB in HEK293T cells and subsequent purification (**Supplementary Fig. 19**), the sensor was assessed for its ability to detect three different anti-RBD antibodies: a recently discovered neutralizing antibody (47D11) and a commercially available ELISA pair (D001 and D003). **Fig. 4g** depicts the RAPPID response curves of these antibodies, with LODs of 106 pM, 16 pM and 4 pM for 47D11, D001, and D003, respectively, and a maximal change in emission ratio of 36-fold. As expected, an antibody (49F1) that binds the spike protein outside the RBD^44^, did not induce a sensor response (**Supplementary Fig. 20**). A similar approach based on intensiometric detection of split NanoLuc complementation was recently published as a preprint^55^, but the ratiometric detection enabled by the RAPPID assay is preferred for quantitative measurements, which are important when studying the time-dependence of the antibody response and the correlation between antibody titers and immunological protection^56,57^. Collectively, these results illustrate that the RAPPID platform is highly versatile and can be reengineered to suit a variety of applications related to antibody detection, including quantitative measurements of antibodies that bind discontinuous epitopes, therapeutic antibodies and their ADAs, and anti-SARS-CoV-2 antibodies.

## Conclusion and discussion

Protein-based biosensors that generate a luminescent signal upon binding of a biomolecular target are attractive bioanalytical tools because they 1) enable highly sensitive detection based on enzymatic amplification, and 2) do not require external excitation, which renders them ideal for direct homogeneous detection in complex samples. In this study, we have established a new comprehensive sensor platform based on antibody-guided split NanoLuc complementation combined with a robust ratiometric output. The RAPPID platform is highly modular and broadly applicable as it allows the use of commercially available monoclonal antibodies as recognition elements. The required protein G-split NanoLuc fusion proteins are easily expressed in *E. coli* and obtained using standard affinity purification, while antibody conjugation is based on a one-hour illumination protocol that does not require expert laboratory equipment. Performing a RAPPID assay entails single-step mixing of the target analyte with the two antibody-luciferase components and the calibrator luciferase, after which NanoLuc substrate is added to initiate signal generation. The ratiometric readout can be recorded in minutes as a blue-to-green color change using a standard digital camera.

Combining these design aspects allowed the formulation of a straightforward workflow for developing in-solution immunoassays with low pM sensitivity, a large dynamic range and excellent reproducibility for any relevant target, depending only on the availability of a suitable pair of antibodies. Sensors for traditional biomarkers such as cardiac troponin I and C-reactive protein, as well as newly emerging targets such as the SARS-CoV-2 virus, could be developed within days after antibody arrival. Depending on the affinity of the antibodies and the target architecture, robust changes in emission ratio ranging from 5 (for ADAs) to 25 were routinely obtained, which is substantially higher than typical BRET-based biosensors. Moreover, the sensor’s responsive range can be rationally adjusted to a specific application by varying the concentration of the antibody conjugates and the concentration of the calibrator luciferase. The platform is versatile by design, which allowed us to create multi-faceted detection assays, for example for measuring both therapeutic antibodies and their anti-drug-antibodies, as well as both SARS-CoV-2 antigens and antibodies. Furthermore, strategic selection of the antibodies can facilitate elucidation of structural features of the target analyte, which could aid in distinguishing between functional and non-functional molecular configurations, for instance when considering the oligomerization state of proteins.

An important aspect of the RAPPID platform is direct in-sample calibration by addition of a calibrator luciferase, that contains an identical luciferase domain but with a spectrally distinct emission profile. Although the ratiometric response is smaller than the intensiometric response, the ratiometric signal is stable over time and less susceptible to experimental variation. We envision that the use of a calibrator luciferase can be applied as a general strategy to translate intensiometric luminescent assays into robust ratiometric assays with a time-independent and quantitative assay output. As such, other recently developed intensiometric sensors based on split luciferase complementation could benefit from a calibrator luciferase to further enhance their potential in a clinically relevant environment^21,55^. Furthermore, the concept can be extended by using different acceptor domains, such as quantum dots, organic fluorophores or fluorescent protein domains, with a larger spectral separation from the blue light-emitting NanoLuc.

In conclusion, RAPPID represents a highly attractive assay format for point-of-care diagnostics, as it combines the analytical performance of a laboratory-based immunoassay such as ELISA with the speed and ease-of-use of lateral flow immunoassays. To allow further development for point-of-care diagnostic applications, RAPPID could be integrated in cheap and simple paper- or thread-based analytical devices or microfluidic cartridges, while dedicated readers could be developed to further increase sensitivity. The availability of such integrated devices and the ability to develop a RAPPID assay for a new target analyte only within days, makes RAPPID an attractive immunoassay format for a broad spectrum of applications, including rapid screening methods, therapeutic antibody drug monitoring, and the rapid detection of infectious diseases to be better prepared for future pandemics.

## Methods

### Cloning

The pET28a(+) vectors containing DNA encoding for Gx-SB, Gx-LB, mNG-NL and SUMO-TNFα-SB were ordered from GenScript. The sequence of mNG-NL was reported by Suzuki et al^32^, and a His-tag at the N-terminus and a Strep-tag at the C-terminus were included to facilitate purification^26^. Site-directed mutagenesis to change SB sequences was carried out using the QuikChange Lightning Site-Directed Mutagenesis kit (Agilent technologies) and specifically designed primers.

The receptor binding domain (RBD) of the SARS-CoV-2 Spike glycoprotein (produced under HHSN272201400008C and obtained through BEI Resources, NIAID, NIH: Vector pCAGGS Containing the SARS-Related Coronavirus 2, Wuhan-Hu-1 Spike Glycoprotein Receptor Binding Domain (RBD), NR-52309) was genetically fused to either the LB or SB via a semi-flexible linker and subsequently cloned into the pHR-CMV-TetO2_3C-mVenus-Twin-Strep plasmid, a gift from A. Radu Aricescu (Addgene plasmid #113891)^58^, by means of restriction-ligation approach.

All cloning and mutagenesis results were confirmed by Sanger sequencing (BaseClear). The DNA and amino acid sequences of the fusion proteins are listed in supplementary information (**Supplementary Fig. 21-26**).

### Protein expression

The pET28a plasmid encoding for either Gx-LB or Gx-SB was co-transformed into *E. coli* BL21 (DE3) together with a pEVOL vector encoding a tRNA/tRNA synthetase pair in order to incorporate para-benzoyl-phenylalanine (pBPA)^27^. The pEVOL vector was a gift from Peter Schultz (Addgene plasmid # 31186). Cells were cultured in 2YT medium (16 g peptone, 5 g NaCl, 10 g yeast extract per liter) containing 30 μg/mL kanamycin and 25 μg/mL chloramphenicol. Protein expression was induced using 0.1 mM IPTG and 0.2% arabinose in the presence of 1 mM pBPA (Bachem, F-2800.0001). Following overnight expression at 20 °C, cells were harvested by centrifugation at 10,000 g for 10 min and lysed using the Bugbuster reagent (Novagen) and Benzonase (Novagen). Proteins were purified using Ni^2+^ affinity chromatography followed by Strep-Tactin purification according to the manufacturer’s instructions. Correct incorporation of pBPA was confirmed by Q-ToF LC-MS (**Supplementary Fig. 7**).

The pET28a plasmid encoding mNG-NL fusion protein was transformed into *E. coli* BL21 (DE3). Cells were cultured in LB medium (10 g peptone, 10 g NaCL, 5 g yeast extract per liter) containing 30 μg/mL kanamycin. Protein expression was induced using 0.1 mM IPTG at 20 °C overnight. The harvested cells were lysed using the Bugbuster reagent and Benzonase. Proteins were purified using Ni^2+^ affinity chromatography and Strep-Tactin purification.

The SARS-COV-2 spike trimer protein was expressed in HEK-293T cells with a C-terminal trimerization motif and Strep-tag using the pCAGGS expression plasmid, and purified from the culture supernatant by Strep-Tactin chromatography as described previously^44^.

The pET28a plasmid encoding SUMO-TNFα-SB fusion protein was transformed into *E. coli* BL21 (DE3). Cells were cultured as described above. The harvested cells were resuspended in Tris-HCl Buffer (40 mM, pH 8.0, containing 125 mM NaCl and 5 mM imidazole) and Benzonase. Cell disruption was then performed by ultrasonication on ice with 5 pulses of 20 s and an amplitude of 50%. After centrifugation, proteins were purified from the supernatant at 4 °C, using Ni^2+^ affinity chromatography. The His-SUMO-tag was cleaved from TNFα-SB by adding 1:1000 (v/v) SUMO protease into the protein solution which was further dialyzed in Tris-HCl buffer (20 mM, pH 8.0, containing 150 mM NaCl and 1 mM DTT) overnight, at 4 °C. Cleaved TNFα-SB was obtained using Ni^2+^ affinity chromatography to remove the cleaved His-SUMO-tag.

For lentiviral production of the two vectors, encoding either the RBD-LB or RBD-SB transgene, HEK293T cells (ATCC® CRL-3216™) were transiently co-transfected with the abovementioned transfer vectors, as well as the VSV-G envelope (pMD2.G) and packaging plasmids (pCMVR8.74), as previously described^58^. Plasmids pMD2.G and pCMVR8.74 were gifts from Didier Trono (Addgene plasmid #12259 and Addgene plasmid #22036, respectively). HEK293T cells were then infected in order to establish polyclonal stable cell lines via the integration of the lentiviral genetic elements into the cell genome. Cells were routinely cultured under standard culturing conditions (37°C, 5% CO2, 95% air, and 95% relative humidity) in Dulbecco’s Modified Eagle Medium (DMEM 21885-025, ThermoFisher), supplemented with 10% Fetal Bovine Serum (FBS). For cell line-maintenance, cells were split at a ratio of 1:5-1:10, upon reaching confluency. Protein expression was induced by the addition of 1 μg/mL doxycycline (D3447, Merck) in DMEM, supplemented with 2% FBS. Successful transduction of cells was monitored by fluorescence imaging of the reporter protein mVenus, ensuing induction. Protein-containing medium was harvested 4-5 days following induction and protein was purified by Strep-Tactin affinity chromatography.

All the purified proteins were snap-frozen and stored at -80 °C until use.

### Photoconjugation

Cardiac Troponin I antibodies 19C7 (4T21) and 4C2 (4T21), and C-reactive protein antibodies C6 (4C28cc) and C135 (4C28) were obtained from Hytest. Therapeutic antibodies cetuximab and infliximab were obtained via the Catherina hospital pharmacy in Eindhoven and Maxima Medisch Centrum pharmacy in Veldhoven, the Netherlands, respectively. Adalimumab (A1048-100) was obtained from Gentaur. Anti-adalimumab/TNFα monoclonal antibody (HCA207) and anti-infliximab (HCA213) were obtained from BioRad. Anti-SARS-COV-2 antibodies 47D11 and 49F1 were produced in HEK-293T cells and purified using Protein-A affinity chromatography as described previously^44^. Commercially available anti-SARS-COV-2 D001, D002, D003 and D004 were ordered from Sino Biological. Photoconjugation reactions were performed using a Promed UVL-30 UV light source (4×9 watt). Samples containing antibody and Gx-LB or Gx-SB in PBS buffer (pH7.4) were illuminated with 365 nm light for 30 to 180 minutes. The samples were kept on ice during photoconjugation. If necessary, the photoconjugated products were further purified using Ni-NTA spin columns (ThermoFisher) followed by PD G10 desalting columns (GE Health) according to the manufacturer’s instructions. Antibody-conjugates were stored at 4 °C or -80°C until use.

### Luminescent assays

Cardiac Troponin I (8T53) and C-reactive protein (8C72) were purchased from Hytest. Anti-cetuximab (HCA221), anti-infliximab (HCA213G) and anti-adalimumab (HCA205) were ordered from BioRad. Similarly, SARS-COV-2 spike monomer was expressed with a C-terminal Strep-tag and purified by Strep-Tactin chromatography^44^. Intensiometric assays were performed at sensor protein concentrations of 0.1-10 nM in a total volume of 20 µL PBS buffer (pH 7.4, 0.1% (w/v) BSA) in PerkinElmer flat white 384-well Optiplate. Measurements in diluted blood plasma were performed by preparing the analytes in pooled human blood plasma and sensor proteins in PBS buffer (pH 7.4, 0.1% (w/v) BSA). After incubation of sensor proteins and analytes for 5-60 minutes, NanoGlo substrate (Promega, N1110) was added at a final dilution of 250 to 1000-fold. Luminescence intensity was recorded on a Tecan Spark 10M plate reader with an integration time of 100 ms. For ratiometric assays, an appropriate amount (0.5-50 pM) of calibrator mNG-NL was added into the samples and luminescence spectra were recorded between 398 nm and 653 nm with a step size of 15 nm, a bandwidth of 25 nm and an integration time of 1000 ms. The blue/green ratio was calculated by dividing bioluminescent emission at 458 nm by emission at 518 nm. LOD and 95% confidence interval of the LOD were calculated by linear regression of the response related to the analyte concentration for a limited range of concentrations (for an overview, see **Supplementary Table 1**).

The luminescence signal was also recorded by using a SONY DSC-RX100 digital camera. The plate was placed into a Styrofoam box to exclude the surrounding light. The pictures were taken through a hole in the box using the digital camera with exposure times of 10-30 s, F value of 1.8 and ISO value of 1600-3200.

## Data availability

The data that support the plots within this paper and other findings of this study are available from the corresponding author upon reasonable request.

## Code availability

Custom-written code for the computer models and simulations that support the experimental findings in this study is available from the corresponding author upon reasonable request.

## Acknowledgements

We thank Dr. Leo IJzendoorn and the members of the T.E.S.T. student team for help with the anti-adalumimab assay, Prof. Frank Grosveld (Erasmus MC, Rotterdam, The Netherlands) for providing access to the 47D11 and 49F1 antibodies, Dr. Maarten Broeren and Dr. Luc Derijks (Máxima Medical Center, Veldhoven, The Netherlands) for useful discussions regarding TNFα-binding therapeutic antibodies. We thank Dr. Simone Wouters for help with the expression of the calibrator luciferase, and Pim de Vink for initial help with the thermodynamic model. This work was supported by the European Research Council, ERC starting grant (ERC-2011-StG 280255) and an ERC proof of concept grant (ERC-2016-PoC 755471), funding by the TU/e COVID19 University Fund, grants from the Netherlands Organisation for Scientific Research, NWO-Take-off-1 grant (NWO, 17820), and RAAK.PRO Printing makes sense (RAAK.PRO02.066).

## Author contributions

Y.N., B.R., and E.v.A designed the study, performed experiments, developed the thermodynamic model, analysed the data, and wrote the manuscript. E.H. developed and characterized RAPPID assays for TNFα antibodies and ADAs. L.B. and contributed to the spike protein assays. A.M.P. developed the expression of the RBD-NL fusion proteins, B.T., C.V. and S.R. contributed to initial experiments for various RAPPID assays, R.A. supervised early experiments for this work that provided proof of principle, W.L. expressed and purified spike proteins and antibodies. B.B. and F.v.K. supervised the spike protein work and provided feedback on the manuscript. T.d.G. supervised the development of the thermodynamic model and provided critical feedback on the manuscript. M.M. conceived, designed, and supervised the study, analysed the data, and wrote the manuscript. All authors discussed the results and commented on the manuscript.

